# Activation of cytoplasmic dynein through microtubule crossbridging

**DOI:** 10.1101/2020.04.13.038950

**Authors:** Manas Chakraborty, Algirdas Toleikis, Nida Siddiqui, Robert A. Cross, Anne Straube

## Abstract

Cytoplasmic dynein is the main microtubule-minus-end-directed transporter of cellular cargo in animal cells [1, 2]. Cytoplasmic dynein also functions in the organisation and positioning of mitotic spindles [3, 4] and the formation of ordered microtubule arrays in neurons and muscle [5, 6]. Activation of the motor for cargo transport is thought to require formation of a complex with dynactin and a cargo adapter [7-10]. Here we show that recombinant human dynein can crossbridge neighbouring microtubules and can be activated by this crossbridging to slide and polarity-sort microtubule bundles. While single molecules of human dynein are predominantly static or diffusive on single microtubules, they walk processively for 1.5 μm on average along the microtubule bundles they form. Speed and force output of dynein are doubled on bundles compared to single microtubules, indicating that the crossbridging dynein steps equivalently on two microtubules. Our data are consistent with a model of autoactivation through the physical separation of dynein motor domains when crossbridging two microtubules. This enables cytoplasmic dynein to function effectively as a microtubule organiser and transporter without needing to first form a complex with dynactin and a cargo adapter.

## Results

Mammalian cells express three different dyneins: axonemal dynein is integral to the structure and force generation of motile cilia and flagella, cytoplasmic dynein 2 mediates retrograde intraflagellar transport, while cytoplasmic dynein 1 (hereafter referred to as dynein) is the major minus-end directed transporter in cells [1, 2, 11]. Dynein cargoes include nuclei and other organelles in addition to recycling vesicles, mRNAs, viruses and other microtubules [12]. Indeed, dynein is a major microtubule organiser implicated in removing minus end out microtubules from axons [5, 13], forming paraxial microtubule arrays in muscle cells [6], focussing the poles of mitotic spindles [3] and balancing centrosome separation forces [14, 15]. Human dynein is a complex of two heavy chains (DHC), two intermediate chains (DIC), two light intermediate chains (DLIC), and three light chain dimers (LC8, Tctex, and Robl) [8, 16-18]. Each heavy chain contains a motor domain related to AAA+ ATPases with a microtubule binding domain at the end of a stalk that changes conformation during the ATP hydrolysis cycle, enabling the motor to step along microtubules [18-20]. However, the human dynein complex is a poorly processive motor and only generates forces of up to 1 or 2 pN [21-23]. To move processively, dynein forms a tripartite complex with dynactin and a cargo adaptor protein such as BICD2, BICDR1 or HOOK3 [8-10, 24, 25]. These dynein-dynactin-adaptor complexes move faster and produce significantly larger forces (up to 6 pN) than dynein alone [22, 25]. This ensures activation of dynein upon loading to a cargo. However, it is unclear how dynein is activated when organizing microtubule arrays and performing microtubule-microtubule sliding. A previous study suggested that dynein slides microtubules relative to each other by walking with its two motor domains on two separate microtubules [26]. Thus, the second microtubule is not attached to dynein’s tail as other cargoes, potentially negating the requirement for activation via cargo adaptor binding.

To test this idea, we purified native cytoplasmic dynein from porcine brain and recombinant human dynein from insect cells (Supplementary Fig. S1). We immobilised dynein onto glass coverslips using antibodies specific to dynein intermediate chain to ensure that motor domains are oriented towards microtubules in gliding assays (Fig. 1A). We observed that microtubule structures increased in length over the course of the experiment and formed bundles (Fig. 1B). Both when gliding on pig brain dynein or recombinant human dynein, the average length of microtubule structures had increased by about 50% after 40 minutes (Fig. 1C). In contrast, gliding on kinesin-1 (*Drosophila* kinesin heavy chain) under identical conditions did not result in a significant change of microtubule length (Fig. 1C-E). The dynein-dependent length increase occurs because individual microtubules were being joined by surface-anchored dynein to form microtubule bundles or chains (Fig. 1F, Supplementary Movie 1), presumably because they are being pulled together by the two motor domains of dynein dimers.

**Figure 1:**
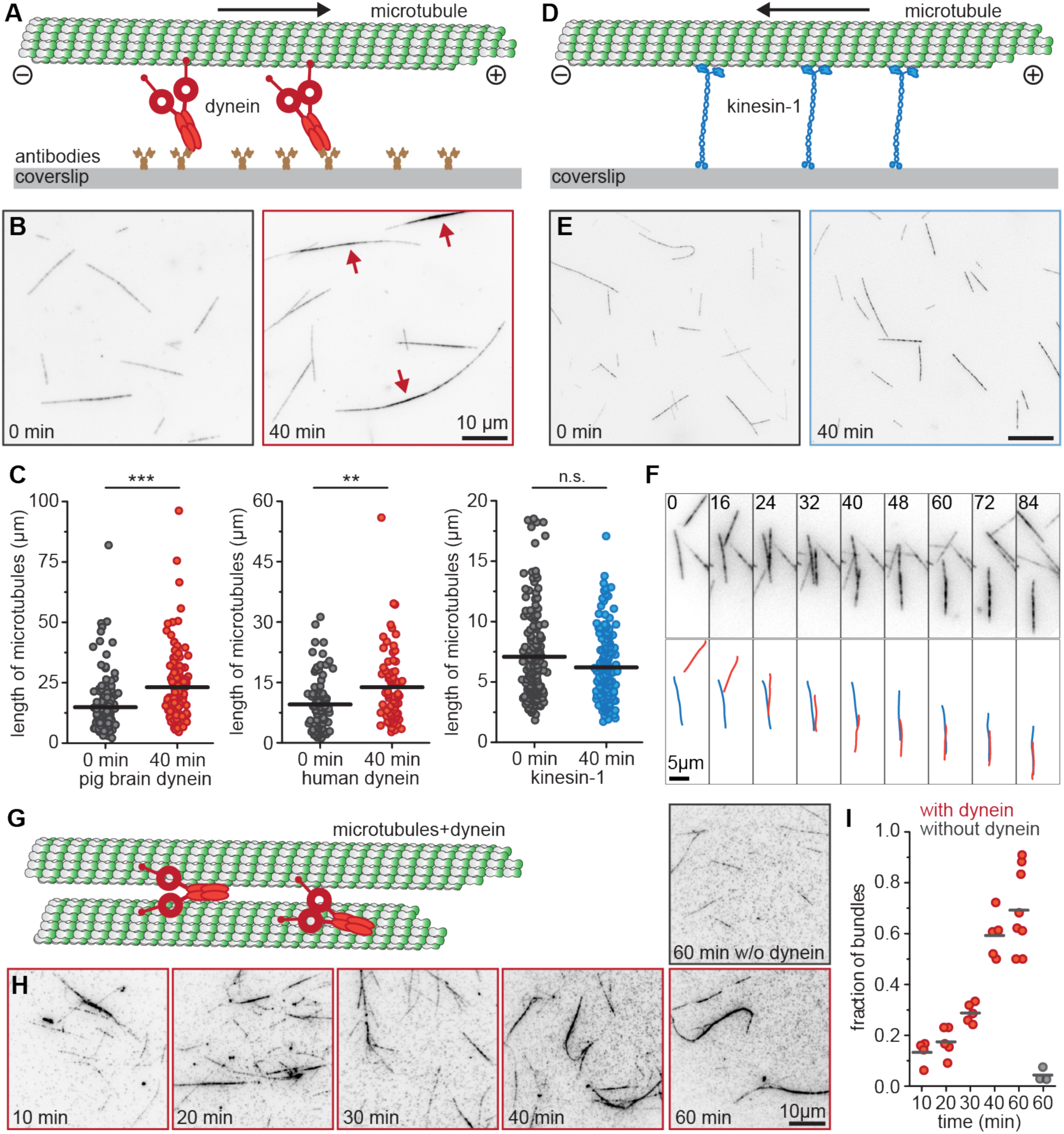
Dynein bundles microtubules. **(A)** Schematic of the microtubule bundling assay by surface anchored dynein. **(B)** Representative fields of view from gliding assays with immobilised dynein at start of assay and after 40 minutes. Note the formation of long bundles or chains (red arrows). Scale bar 10 µm. **(C)** Lengths of microtubule structures are shown for gliding assays with brain dynein, recombinant human dynein and Drosophila kinesin-1 at 0 minute and 40 minute. The bar indicates the mean. n = 159,129,88,74,170,166 respectively pooled from 3 independent experiments. *** p<0.0001, ** p<0.001, n.s. p>0.5 (Mann-Whitney U-test). **(D)** Schematic of microtubule gliding assay with surface anchored kinesin. **(E)** Representative fields of view from gliding assays with immobilised kinesin-1 at start of assay and after 40 minutes. Scale bar 10 µm. **(F)** Time series showing how microtubules join into chains and bundles during gliding assays on a dynein-coated surface. Times in seconds are indicated in top left corner. Scale bar 5 µm. **(G-H)** Bundling of microtubules with recombinant human dynein in solution. Representative images after different incubation times are shown. Scale bar 10 µm. **(I)** Fraction of bundled microtubules as a function of time incubated with recombinant human dynein. Data show individual data points and mean from 3-8 independent experiments with >100 microtubules analysed for each dataset. Control data are 60 minutes incubation without dynein.

We next investigated microtubule bundling in the presence of fluorescently-labelled, recombinant dynein in solution (Fig. 1G). Consistently with our gliding assays, microtubules were increasingly bundled by dynein over the course of one hour (Fig. 1H-I). To determine the polarity of the microtubules within the bundles, we used two complementary approaches. First, we added dynein-mediated bundles into a chamber containing surface-immobilised kinesin-1 (Fig. 2A). The rationale was that kinesin-1 will split microtubule bundles and by observing the direction of motion of component microtubules we can directly determine the polarity of microtubules in the bundles. Almost all microtubule bundles were split apart on the kinesin surface (Fig. 2B, and Supplementary Movie 2). On average bundles consisted of 3.8±0.3 microtubules and the vast majority of these bundles (>85%) released microtubules that moved in the same direction as the original bundle (Fig. 2C), thus dynein forms microtubule bundles that are predominantly parallel. As an alternative approach, we determined microtubule orientation by analysing the motility of dynein molecules in the bundled microtubule structures (Fig. 2D-E). Microtubule polarity in the bundles was inferred from the ratio of dynein runs in the same direction as the majority of dynein runs to the total number of dynein runs. Therefore, a polarity index of 0.5 indicates an antiparallel microtubule bundle, while 1 indicates a parallel bundle. We found that 75% of the bundles we examined had a polarity index larger than 0.8 (Fig. 2F). Interestingly, the microtubule bundles formed after shorter incubation times with dynein have a significantly lower average polarity index than bundles formed after prolonged incubation with dynein (10-30 minutes: 0.80 ± 0.03 (mean ± SEM), n = 34 bundles; 40-60 minutes: 0.94 ± 0.02, n = 70 bundles; Wilcoxon ranksum test p = 2.8•10^−4^). This suggests that dynein might first bundle microtubules in a random orientation and subsequently sort them. When observing the motion of bundled microtubules relative to each other, we observe that parallel bundles do not to show any sliding motion, but a sizable portion (24%) of antiparallel bundles (bundling index <1) slide apart at an average velocity of 37 ± 8 nm s^-1^ (Fig. 3A-C). Any microtubules driven apart usually detach and float away, while occasionally flipping over into a parallel configuration. This suggests that bundles form randomly in the presence of dynein and antiparallel bundles are specifically removed by microtubule-microtubule sliding, resulting in an enrichment of parallel bundles. Thus, dynein exhibits a similar polarity-sorted bunding activity as the minus-end directed kinesins Klp2 and Ncd [27, 28]. However, with the difference that kinesin-14s have an additional ATP-independent microtubule binding site in the tail, while dynein is thought to move with each motor domain on two separate microtubules [26]. To obtain direct evidence for this idea, we simultaneously observed sliding microtubules and labelled dynein. We expected and found a mixture of behaviours amongst dynein molecules, consistent with 5 different motility states: molecules are static on either the track or the transport microtubule, molecules that move along only one of the microtubules and molecules that generate force on both microtubules and therefore move relative to the substrate at a speed about half of the microtubule sliding speed (Fig. 3D). We observed a significant fraction of molecules moving at intermediate speeds (Fig. 3D-E, Supplementary Movie 3), supporting the notion that a dynein molecule crosslinks two microtubules with its motor domains and walks on both of them. While analysing microtubule polarity in bundles, we noticed that a significant fraction of single dynein motors moved processively. This was surprising as full length human dynein has been reported to be mainly static, occasionally diffusive and at most poorly processive [8, 10, 16, 29]. Comparing the motility of dynein molecules on single microtubules and bundles from the same field of view, suggested that dynein resides for about the same time on single and bundled microtubules, only rarely undertaking directional runs on single microtubules, while these were frequently observed on microtubule bundles (Fig. 4A-C). Dynein molecules exhibited processive motion of 1.27 ± 0.06 µm average run length on bundles, but only managed to move 0.23 ± 0.04 μm on average on single microtubules (Fig. 3C). If we disregard all static dynein molecules, i.e. those with a displacement of less than 2 pixels (162 nm), the average run length of dynein on bundles (1.50 ± 0.06 µm) is still more than twice as long as for runs on single microtubules (0.64 ± 0.10 µm). Thus even dynein molecules that are not fully autoinhibited, rarely complete runs of 1 µm or more on single microtubules in line with previous publications [29]. In addition, dynein moved much faster on bundles than on single microtubules (Fig. 3D) (134 ± 6 vs 25 ± 4 nm s^-1^; mean ± SEM, n = 576, 188 molecules), and speed more than doubled even if only considering non-static motors (159 ± 7 vs 68 ± 12 nm s^-1^; mean ± SEM, n = 482, 59 molecules). To exclude the possibility that dynein multimers form in bundles and result in the improved processivity, we confirmed that the fluorescence intensity of dynein particles observed on single and bundle microtubules in these experiments is comparable (Supplementary Fig. S2A). Appreciable motor activity is usually only observed for human dynein after activation by forming a ternary complex with dynactin and a cargo adapter. Our observations suggested that crossbridging two microtubules might also activate the motor without the need for additional cofactors. Since dynein has previously been shown to generate higher forces when activated by dynactin and a cargo adapter, we next compared the force output of dynein complexes on single and bundled microtubules using an optical trap. Dynein was immobilised on 560 nm polystyrene beads at a dilution resulting in less than 25% of beads moving when brought near surface-immobilised microtubule bundles (Fig. 4E-G). This ensured that 90% of observed bead movements are due to a single motor engaging with the microtubules. As the probability of engaging with a microtubule bundle is higher than a single microtubule, we observed a higher frequency of runs on bundles than on single microtubules (Fig. 4H). In line with previous work [23, 29-31], we find that recombinant human dynein produces brief movements with an average stall force of 0.89±0.06 pN (mean±SEM, n=105 runs from 7 experiments) on single microtubules (Fig. 4I-L). However, on bundles, we regularly observe dynein to generate forces of several pN with an average stall force of 1.65±0.07 pN (mean±SEM, n=236 runs from 3 experiments) (Fig. 4J-L, Supplementary Figure S3). Force-velocity data suggest that dynein can sustain significant directional motion against 6 pN force when moving on bundles, but only move against a 3 pN load on single microtubules (Fig. 4M).

**Figure 2:**
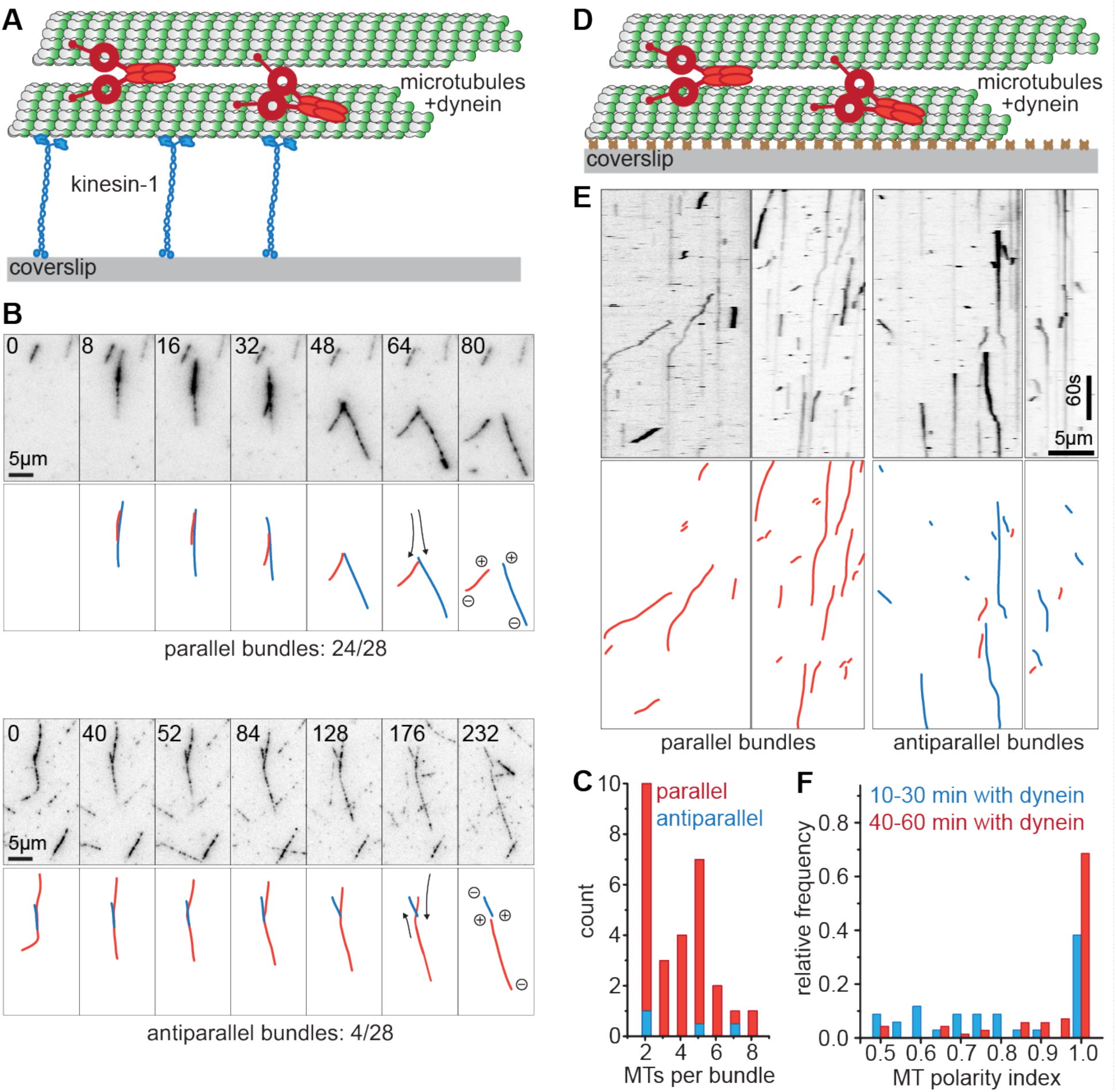
Dynein forms predominantly parallel bundles. **(A)** Schematic of the microtubule polarity analysis of dynein-formed bundles using surface anchored kinesin. **(B)** Representative time-lapse images from gliding assays with immobilised Kinesin-1 and dynein-mediated microtubule bundles. Relative time in seconds is indicate in the top left corner. Note a moving bundle splits and component microtubule (red and blue) move either in the same direction (top panel), or in opposite direction (bottom panel). Scale bar 5 µm. **(C)** Stacked histogram of the number of microtubules in a bundle and their relative orientation. Red bars indicate that all component microtubules were parallel while blue bars indicate bundles with at least one component microtubule moving in the opposite direction. n = 28 microtubule bundles from 3 independent experiments. **(D)** Schematic of the microtubule polarity analysis of dynein-formed bundles using single-molecule TIRF microscopy. **(E)** Representative kymographs showing dynein runs in microtubule bundles. Molecules moving from right to left are highlighted in red, while molecules moving left to right are indicated in blue. Scale bars 5 µm (horizontal), and 60 s (vertical). **(F)** Histogram of microtubule polarity index determined as ratio of parallel dynein runs to total dynein runs for samples incubated 10-30 minute with dynein (blue) or 40-60 minute with dynein (red). n=105 microtubule bundles, from 3 experiments.

**Figure 3:**
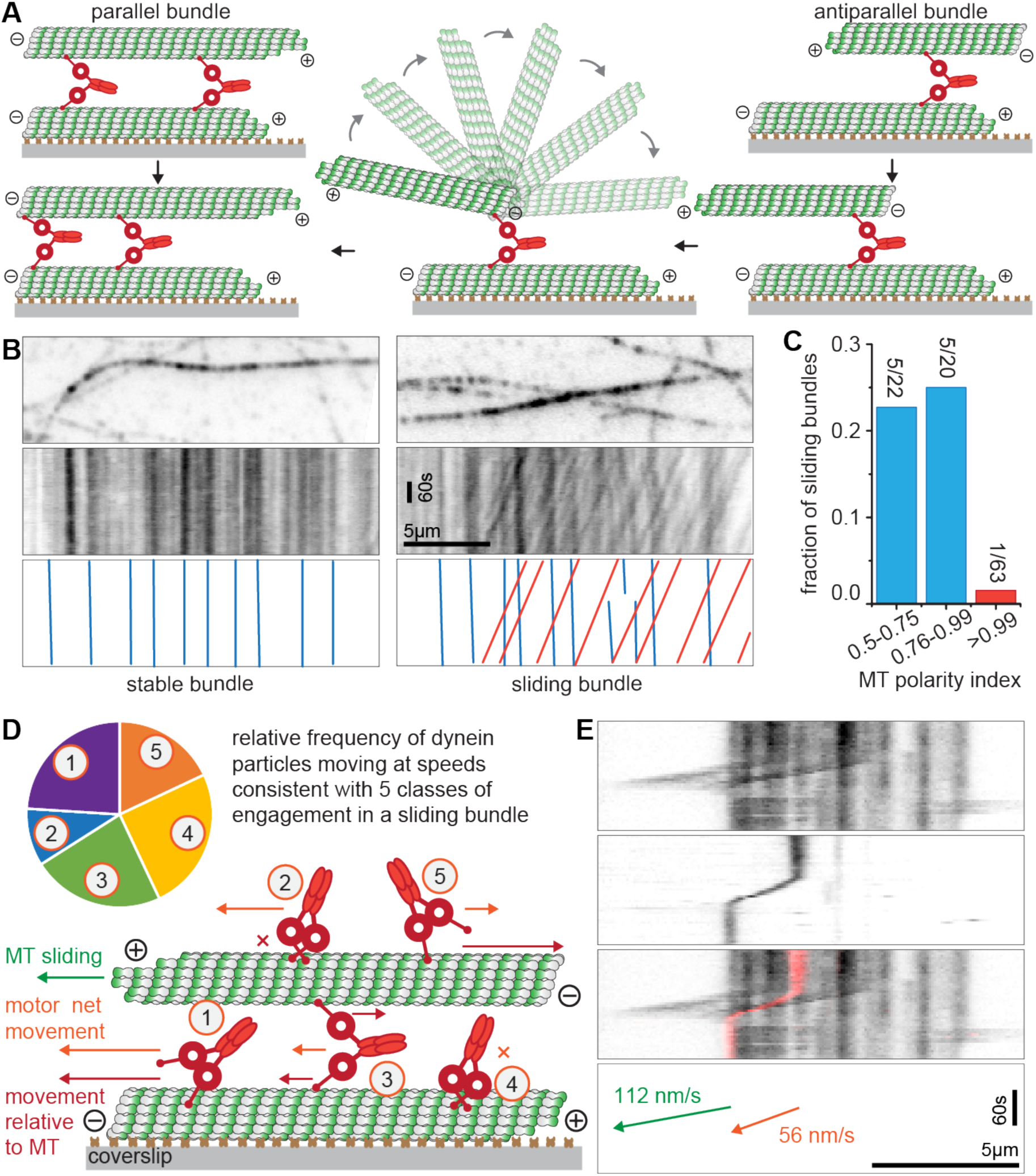
Polarity-sorting of bundles by dynein. **(A)** Schematic of polarity sorting due to preferential sliding of antiparallel-oriented microtubules that either result in direct conversion into a parallel bundle if microtubule swivels around (as shown here) or dissolving the bundle if one microtubule detaches. Parallel bundles don’t move and are therefore enriched. **(B)** Example image and kymographs of microtubule bundles without (left) or with (right) noticeable microtubule-microtubule sliding motion. **(C)** Fraction of sliding bundles with different microtubule polarity index as determined by directionality of dynein runs. Data from n = 105 microtubule bundles from 3 independent experiments. **(D)** Schematic of 5 different classes of dynein engaging with one or both microtubules in an antiparallel bundle and frequency of net speed relative to sliding speed observed that are consistent with each class. (1) Moving on track microtubule (speed faster than microtubule sliding), (2) Static on transport microtubule (same speed as microtubule sliding ± 12%) (3) Force generation on both microtubules (intermediate speed) (4) Static on track microtubule (no movement ± 12%) (5) Moving on transport microtubule (significant movement in opposite direction). n = 100 molecules in 10 sliding microtubule bundles. **(E)** Kymograph of an antiparallel bundle showing microtubules (top panel), dynein (second panel), merge (third panel) and measured speeds (bottom panel). Motility of dynein at half microtubule sliding speed is consistent with walking on both microtubules it crosslinks.

**Figure 4:**
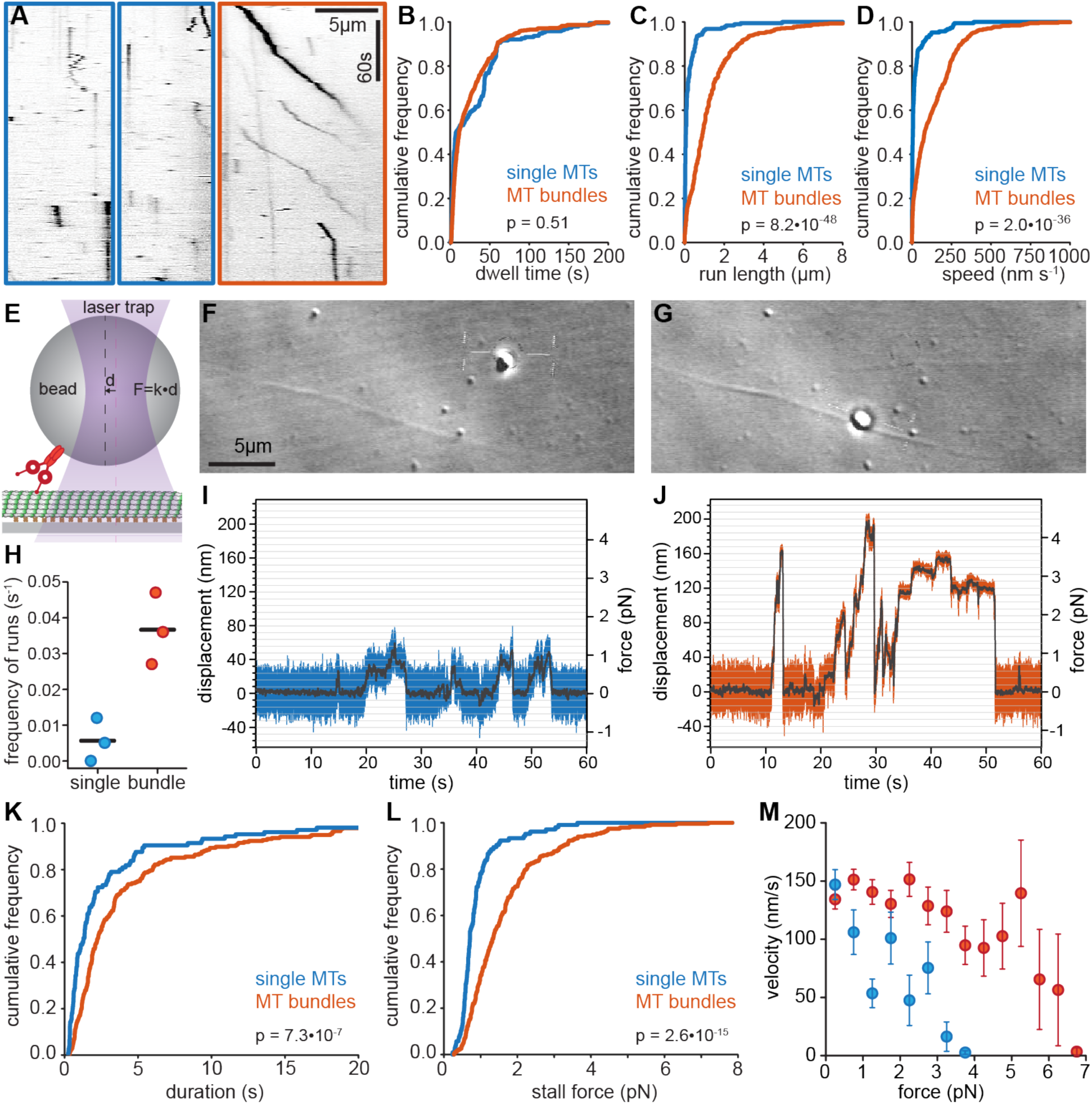
Speed, processivity and force output of human dynein increases on microtubule bundles. **(A)** Representative kymographs showing single molecule behaviour for human dynein on single (blue outline) and bundled microtubules (red outline). **(B-D)** Dwell time, run length and average speed are shown as cumulative distributions for dynein on single and bundled microtubules as indicated. All binding events lasting more than 1s were included. n=188, 576 molecules from 47, 34 microtubules from 3 independent experiments. p value is shown for Wilcoxon rank sum test. **(E)** Schematic representation of single dynein trapping experiment. **(F-G)** Beads are brought close to single microtubules (F) and dynein-mediated microtubule bundles (G) and displacements recorded. Scale bar 5 µm. **(H)** Frequency of runs observed on single versus bundled microtubules under single molecule conditions for all beads that produced at least one run. A run is defined as an event where stall force was sustained with ±10% fluctuation for more than 100ms followed by a sudden force drop to baseline with a displacement of at least 24nm. **(I-J)** Representative traces from single molecule optical trapping experiments of the same bead with dynein on single microtubules (I) or a microtubule bundle (J). Raw displacement and force data at 1000 Hz are shown in colour and smoothed data at 10 Hz are shown in grey. **(K-L)** Cumulative frequency plot of duration of runs (K) and stall force reached (L) by dynein molecules on bundled versus single microtubules. See (H) for definition of run. n=105, 236 events from 7, 3 independent experiments, respectively. p value is shown for Wilcoxon rank sum test. **(M)** Force-velocity relationship for dynein molecules on single (blue) and bundled (red) microtubules extracted from manually identified runs over fixed force windows of 0.5pN. Data show mean ± SEM. Analysis was conducted on 105, 236 runs from 7, 3 independent experiments.

## Discussion

Dynein autoinhibition is thought to involve stacking of the motor domains against each other, trapping of the linker that mediates the powerstroke, and positioning the microtubule domains in opposite orientation [18, 29, 32]. We envisage that microtubule crossbridging activates dynein by stably separating motor domains, in opposition to their intrinsic tendency to autoinhibit and that it is this separation that causes the observed increase in speed, processivity and force generation (Supplementary Fig. S2B). This is consistent with previous observations that dynein autoinhibition can be at least partially overcome by preventing the stacking of the two motor domains. Coupling two dynein motor domains tightly to a short actin filament as spacer doubled run length and speed [29]. For cytoplasmic dynein-2, mutating residues responsible for electrostatic interactions of the stacked motor domains, resulted in about threefold increased speed in gliding assays [32]. Similar interface mutations also increased run length of dynein/dynactin/adapter complexes two-fold [18]. Activation of the dynein holoenzyme by crossbridging within microtubule bundles appears more effective. The mean residence times appear similar for dynein attached to a single microtubule and dynein crossbridging two microtubules in a bundle, suggesting broadly equivalent interactions of each head with the microtubule lattice in the two cases. Together, our data show that dynein holoenzyme is activated by crossbridging microtubules, so that the two motor domains bind different microtubules, and that the resulting ability to slide antiparallel microtubules or move processively on parallel microtubules is robust under load.

Dynein’s involvement in the sliding and polarity sorting of microtubules in cells is well established [3, 5, 6, 13-15, 33]. Interestingly, the dynein co-factor dynactin is not required for all dynein-dependent functions in mitosis [34]. While dynactin is required for localising dynein to the kinetochore and the nuclear envelope, for silencing the spindle assembly checkpoint and for anchoring the centrosome, it is dispensable for pole focussing, for generating contractile forces in the spindle and for chromosome congression [34]. The latter functions primarily involve dynein mediating microtubule-microtubule sliding and moving towards the minus ends of parallel microtubule bundles. Further, the dynein tail, which binds to dynactin, is dispensable for microtubule-microtubule sliding in vitro and in vivo [26]. Nonetheless, other dynein co-factors are required for these cellular functions, especially Lis1 [34], which binds to the motor domain near AAA3 and regulates the coupling between ATP hydrolysis and microtubule binding/release. Lis1 can prolong the engagement of dynein with microtubules under load [23] or facilitate faster stepping depending on whether one or two Lis1 molecules are bound to the motor domain [35]. Thus, the intrinsic ability of dynein holoenzyme to be activated by microtubule crossbridging is likely in many cases to be further regulated and further work will be required to elucidate the interplay of the various regulatory mechanisms.

We anticipate that microtubule crossbridging autoactivates the dynein holoenzyme by stabilising the physical separation of the motor domains, in opposition to autoinhibition via stacking of the AAA domains, stabilised by their mutual electrostatic attraction. Mutants that prevent motor domain stacking should thus be hyperactive, especially for those functions that involve microtubule sliding and processive motion towards the minus end of bundles. Indeed, such hyperactive mutants over-accumulate at spindle poles and result in monopolar spindles [18], presumably because the balance between dynein-mediated contractile forces now exceeds the extensile forces generated by Kif11 and Kif15. As a number of mutations causing neuromuscular disease reside in the AAA interface [18, 36], the balanced regulation of dynein activity is likely to be of key importance also in post-mitotic cells such as neurons and muscle, in which dynein functions as a key microtubule organiser and transporter.

## Acknowledgements

We thank Andrew Carter (MRC-LMB Cambridge) for Multibac plasmids and DH10BacY for the expression of recombinant human dynein in insect cells and the sharing of purification protocols for dynein and TEV protease. This work was funded by a Wellcome Investigator Award in Science (200870/Z/16/Z) to A.S. and a Senior Wellcome Investigator Award (103895/Z/14/Z) to R.A.C.

## Author contributions

MC, RAC and AS conceived the project and designed experiments. AT performed optical trapping experiments and analysed data, NS purified kinesin-1 and analysed data, MC performed all other experiments and analysed data. MC and AS wrote the manuscript with contributions from all authors.

## Figures and Figure Legends

**Supplementary Figure S1:**
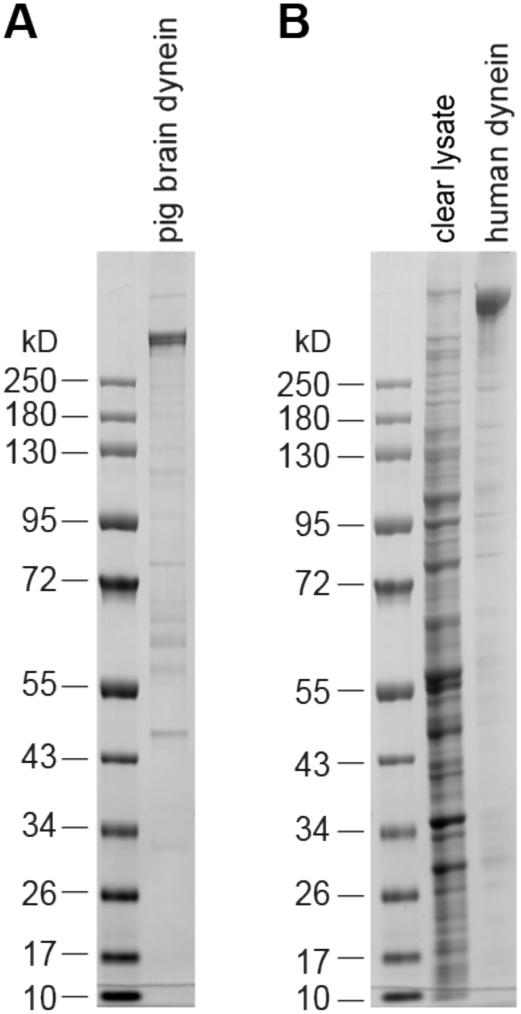
Purified dynein. **(A)** Purified dynein from porcine brain. **(B)** Insect cell lysate expressing recombinant human dynein and the purified complex. Molecular weight markers in kD as indicated.

**Supplementary Figure S2:**
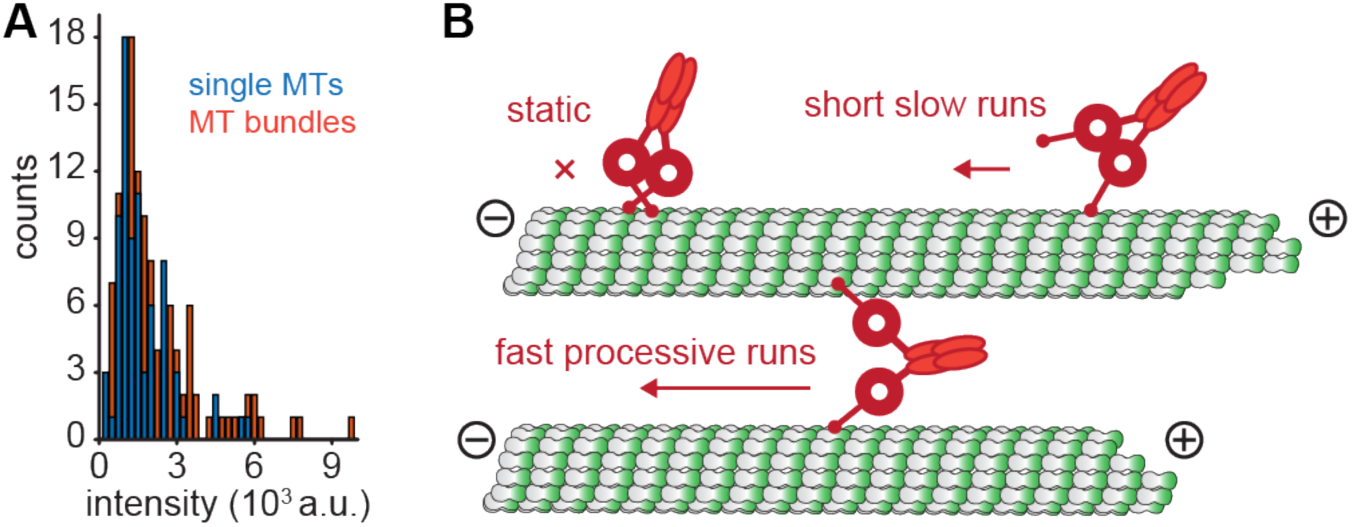
Model of substrate-activated dynein. **(A)** Intensity distributions of dynein particles on single microtubules and microtubule bundles. n=79, 127 molecules from 4 experiments. p=0.057, t-test. **(B)** Model showing substrate-dependent activation of human cytoplasmic dynein on microtubule bundles. The majority of dynein in solution is in an autoinhibited form. Autoinhibited dynein adopts a phi conformation with microtubule-binding domains pointing in opposite directions and motor domains in contact. This allows dynein to bind with one motor domain at a time and results in static or diffusive interactions. Occasional opening of the motor allows slow directional runs on single microtubules as the probability of the motor domains interacting with each other is high. Interaction with two separate microtubules in a bundle allows dynein to undertake processive runs resulting either in the sliding of an antiparallel bundle or fast translocation of dynein towards the minus end of the bundle.

**Supplementary Figure S3:**
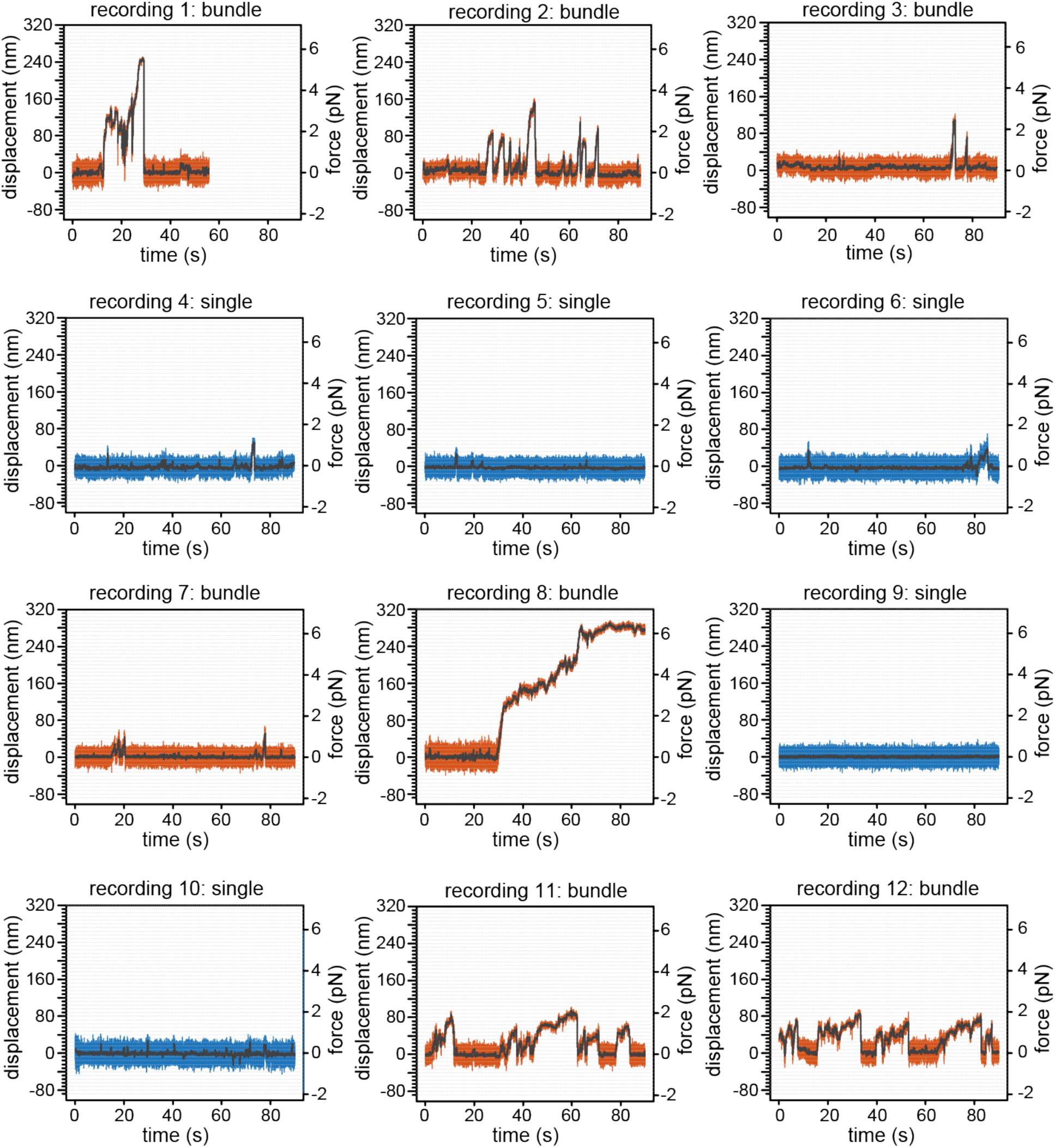
Recordings from optical trapping experiments. Consecutive recordings using the same bead on single microtubules (blue traces) and microtubule bundles (orange traces). Raw displacement and force data at 1000 Hz are shown in colour and smoothed data at 10 Hz are shown in grey.

## METHODS

### Protein purification

Dynein from porcine brain was purified using a microtubule affinity-based purification protocol as described previously [37]. Briefly, ∼75 g porcine brain white matter was blended in cold extraction buffer (0.05 M PIPES-NaOH, 0.05 M HEPES, pH 7.0, containing 2 mM MgCl_2_, 1 mM EDTA, 1 mM phenylmethylsulfonyl fluoride (PMSF, #MB2001, Melford), 10 µg ml^-1^ leupeptin (#L2884, Sigma), 10 µg ml^-1^ tosyl arginine methyl ester (TAME, #283096 Sigma), 1 µg ml^-1^ pepstatin A (#P5318, Sigma), and 1 mM DTT (#MB1015 Melford)). The homogenate was centrifuged at 24000 × *g* (SLA1500 rotor, Thermo Fisher Scientific) for 30 minutes at 4 °C to collect the low speed extract, which was further centrifuged at 150,000 *× g* (Type 60 Ti rotor, Beckman) for 60 minutes at 4 °C to collect cytosolic extract (CE). Taxol was added to the CE at a final concentration of 20 µM and the mixture was incubated at 37 °C for 12 minutes. The CE was then loaded onto 8 ml prewarmed 7.5% sucrose solution in extraction buffer and microtubules pelleted at 40,000 × *g* (SS34 rotor, Thermo Fisher Scientific) for 30 minutes at 37 °C. The microtubule pellet was resuspended in extraction buffer (20% of CE volume) at RT. Taxol was added to the suspension at a final concentration of 5 µM, which was then incubated at 37 °C for 10 minutes followed by centrifugation at 40,000 × *g* (SS34 rotor) for 30 minutes at 25 °C. The resulting pellet was resuspended in extraction buffer, incubated for 10 minutes with 5 µM taxol and spun at 40,000 × *g* (Type 60 Ti rotor) for 30 minutes at 25 °C. The resulting pellet was resuspended in extraction buffer (10% of CE volume) and supplemented with 3 mM Mg-GTP. Resuspension was done using a 15 ml douncer (#885301, Kontes) with 40 strokes. The suspension was then supplemented with 5 µM taxol followed by incubation for 10 minutes at 37 °C. The microtubule pellet was collected by centrifugation at 40,000 × *g* (Type 60 Ti rotor) as before. The pellet was resuspended in extraction buffer containing 10 mM Mg-ATP (#A9187, Sigma-Aldrich) for 10 minutes at 37 °C. The suspension was then centrifuged at 200,000 × *g* (Type 60 Ti rotor) for 30 minutes at 25 °C to obtain a supernatant enriched with dynein. The ATP extract was further fractionated on a 5-20% sucrose gradient in Tris-KCl buffer (20 mM Tris-HCl, pH 7.6, containing 50 mM KCl, 5 mM MgSO_4_, and 0.5 mM EDTA) supplemented with 1 mM DTT at 125,000 × *g* (SW Ti40 rotor, Beckman) for 16 hours at 4 °C. The gradients were separated into 650 µl fractions, and SDS PAGE gel electrophoresis was used to determine the fraction containing dynein. The fractions containing dynein were pooled, aliquoted, flash frozen and stored in liquid nitrogen. Active brain dynein for gliding assays was further purified immediately prior to undertaking any experiments. To do this, dynein solution was incubated with microtubules and 2 mM Mg-ATP at RT for 20 minutes, followed by centrifugation at 20,000 × *g* for 15 minutes at RT. This removed inactive dynein molecules that pellet together with microtubules. The supernatant enriched in active dynein was collected and placed on ice for immediate use.

Recombinant full-length human dynein was purified from Sf9 insect cells (#EM71104-3, VWR) using the multigene baculovirus expression system [38] as described previously [8] with few modifications. Briefly, pDyn1 encoding human cytoplasmic dynein heavy chain (DYNC1H1) codon-optimized for insect cells with an N-terminal SNAPf tag for fluorescent labelling and a TEV-cleavable His-ZZ tag for purification and pDyn2 encoding DYNC1I2 (DIC2), DYNC1LI2 (DLIC2), DYNLT1 (Tctex1), DYNLL1 (LC8) and DYNLRB1 (Robl1) were recombined using Cre-lox recombinase (#M0298S, New England Biolabs) to obtain pDyn3 which was then transformed into *E. coli* DH10BacY competent cells and plated on LB-Agar plates supplemented with 30 µg ml^-1^ kanamycin (#K4000, Sigma), 7 µg ml^-1^ gentamycin (#G1372, Sigma), 10 µg ml^-1^ tetracycline (#T3258, Sigma), 40 µg ml^-1^ Isopropyl β-D-1-thiogalactopyranoside (IPTG, #MB1008, Melford) and 100 µg ml^-1^ X-Gal (#MB1001, Melford). White colonies (positive transformants) were screened by PCR (see Supplementary Table 1 for oligonucleotides used) to confirm integration into the baculovirus genome and the presence of all expression cassettes. Bacmid DNA was purified from positive colonies using alkaline lysis, and bacmids were used to transfect Sf9 cells at 0.5 × 10^6^ cells ml^-1^ density in SF-900 II media (#10902088, Thermo Fisher Scientific) using Escort IV (#L-3287, Sigma) transfection agent following manufacturer’s guidelines. Cells were incubated for 3-5 days at 27 °C and transfection efficiency was monitored regularly by observing YFP expression level using a fluorescence microscope (DeltaVision, Applied Precision, LLC). At >50% transfection efficiency, 2ml cell suspension (P1 virus) was used to inoculate a fresh 50 ml culture of Sf9 cells at 1.5 × 10^6^ cells ml^-1^ density and grown in an incubator shaker (Thermo Scientific) at 27°C, 120 rpm for 72 hours. Cells were pelleted at 1,500 *× g* for 7 minutes at 4 °C and the supernatant (P2 virus) was used to infect Sf9 cells at 1.5 × 10^6^ cells ml^-1^ density. Expression was followed over 5 days of incubation by monitoring YFP fluorescence as above. Cells were harvested when ∼90% cells were fluorescent (usually 72 hours post infection) by centrifugation (252 *× g*, 20 minutes, 4 °C). Cell pellets were flash frozen and stored at -80 °C. Thawed cell pellets were lysed in 10 ml lysis buffer (50 mM HEPES pH 7.4, 100 mM NaCl, 1 mM DTT (#MB1015, Melford), 0.1 mM ATP (#A30030, Melford), 10% (v/v) glycerol, 2 mM PMSF (#MB2001; Melford), 1x cOmplete protease inhibitor cocktail (#05056489001, Roche) by using a 15 ml douncer and cleared by centrifugation (186,200 *× g*, 1 hour, 4 °C, T865 rotor (#51411, Thermo Fisher Scientific)). Clarified supernatant was then incubated with 0.5 ml IgG Sepharose 6 FastFlow beads (#17096901, GE Healthcare) for 4 hours, transferred into a disposable 5 ml polypropylene column (#29922, Thermo Fisher Scientific) and thoroughly washed with lysis buffer. Fluorescence labelling was done by incubating with 5 nmol SNAP-Surface Alexa Fluor 488 (#S9129S, New England Biolabs) for 1 hour at 4 °C. This step was omitted when preparing unlabelled dynein. Beads were washed in TEV cleavage buffer (50 mM Tris–HCl pH 7.4, 148 mM KAc, 2 mM MgAc, 1 mM EGTA, 10% (v/v) glycerol, 0.1 mM ATP, 1 mM DTT) to remove any unreacted dye and were incubated with 0.2 µM TEV protease (expressed from pRK793 (Addgene plasmid # 8827) and purified from *E. coli*) at 4 °C overnight for on bead cleavage. Dynein was collected, concentrated, and buffer exchanged into GF50 buffer (25 mM HEPES pH 7.4, 50 mM KCl, 1 mM MgCl_2_, 5 mM DTT, 0.1 mM ATP) using 100 kD MWCO concentrator (#Z40167, Amicon Ultracel, Merck-Millipore). Dynein concentration and labeling efficiency was measured using a spectrophotometer, aliquoted into 5 µl portions, flash frozen and stored in liquid nitrogen.

Recombinant Kinesin Heavy Chain from *Drosophila melanogaster* was purified as described previously [39]. Briefly, BL21 DE3 (Invitrogen) were transformed with plasmid pPK113-6H-DHK (accession # AF053733), and the cells were grown at 37 °C until OD_600nm_ reached 0.5. Expression was induced with 0.4 mM Isopropyl β-D-1-thiogalactopyranoside at 15 °C overnight. Cells were harvested by centrifugation (3000 *× g*, 15 minutes, RT). Cell pellets were stored at -80 °C. For purification, pellets were thawed on ice, and cells were lysed by sonication in buffer A (50 mM phosphate buffer pH 7.5, 300 mM NaCl, 10% glycerol, 1 mM MgCl_2_, 0.1 mM ATP, and 40 mM Imidazole). The lysate was clarified by centrifugation (100,000 × *g*, T865 rotor, 30 minutes, 4 °C) and the supernatant was then incubated with 200 µl Ni-NTA (#30230, Qiagen) beads for 1 hour at 4 °C. Beads were washed with buffer A containing 90 mM imidazole. Finally, the protein was eluted with buffer A containing 300 mM imidazole. Unlabelled tubulin was purified from porcine brain using a high molarity PIPES wash protocol [40]. Briefly, a brain homogenate was prepared by blending 400 g porcine brain with equal volume of ice-cold DB buffer (50 mM MES, pH 6.6 with HCl, 1 mM CaCl_2_) at low speed for 30 s. The homogenate was then clarified by centrifugation at 10,000 × *g* (SLA1500 rotor) for 2 hours at 4 °C. The supernatant was mixed with an equal volume (250 ml) of prewarmed HPMB (1 M PIPES, pH 6.9 with KOH, 10 mM MgCl_2_, 20 mM EGTA), 250 ml glycerol, and supplemented with 1.5 mM ATP and 0.5 mM GTP. The mixture was incubated for 60 minutes at 37 °C. Polymerized microtubules were pelleted by centrifugation at 150,000 × *g* at 37 °C for 30 minutes. Pellets were resuspended in ice cold DB buffer in a 40 ml douncer with 40 strokes. The suspension was then kept on ice for 30 minutes to depolymerise microtubules. The suspension was clarified by centrifugation at 70,000 × *g* (T865 rotor) at 4 °C for 30 minutes. The supernatant was mixed with equal volume of prewarmed HPMB (120 ml), 120 ml glycerol, 1.5 mM ATP, 0.5 mM GTP for 30 minutes at 37 °C. Polymerized microtubules were pelleted again at 151,000 × *g* for 30 minutes at 37 °C and then resuspended in 6 ml ice cold MRB80 (80 mM PIPES, pH 6.9 with KOH, 4 mM MgCl_2_, 1 mM EGTA, pH 6.9) and incubated on ice for 30 minutes to depolymerise microtubules. The suspension was clarified by centrifugation at 104,000 × *g* (TLA 100.3 rotor, Beckman) for 30 minutes at 4 °C. The supernatant containing purified tubulin was aliquoted, flash frozen and stored in liquid nitrogen. HiLyte 488 (#TL488M-A, Cytoskeleton Inc.), X-rhodamine (#TL620M-A, Cytoskeleton Inc.), HiLyte 647 (#TL670M-A, Cytoskeleton Inc.), and Biotin-labelled tubulin (#T333P-A, Cytoskeleton Inc.) was purchased from Cytoskeleton Inc.

### Microtubule assembly

GMPCPP-stabilised microtubules were prepared from 3 μl of 12 mg ml^-1^ unlabelled tubulin, 0.4 μl of 5 mg ml^-1^ fluorescently labelled tubulin (HiLyte 647, or X-rhodamine labelled tubulin), and 0.4 μl of 5 mg ml^-1^ biotin-labelled tubulin (optional). Tubulin solutions were mixed with 1 µl 10 mM GMP-CPP (#NU-405S, Jena Bioscience) on ice and incubated for 10 minutes to allow nucleotide exchange. Microtubules were polymerized by incubating the mixture at 37 °C for 30 minutes. 50 µl MRB80 was added to the polymerization mix and microtubules were pelleted by centrifugation (20,238 × *g*, 20 minutes, RT). Microtubule pellets were resuspended in MRB80 supplemented with 10 µM Taxol (#T7402, Sigma-Aldrich) and kept at RT in the dark.

### Microtubule gliding assay

A microscopy perfusion chamber was prepared by attaching a coverslip (Menzel-Glaser, 22 × 22 mm, # 1.5, acid treated, and plasma cleaned) on a glass slide (Menzel-Glasser, Superfrost) by using double sided tape (Scotch). Solutions were exchanged into the perfusion chamber using a pipette and filter paper. The chambers were functionalized by flowing in different reagents and incubating for specified amount of time with intermittent washing (with 30 μl of MRB80) in between solutions. All incubations were done inside a humidified chamber to prevent sample dry up. Kinesin-1 and recombinant dynein were non-specifically absorbed to the surface by flowing in 0.1 μM motor solution in MRB80, supplemented with 0.1 mM Mg-ATP, followed by 30 minutes incubation and washing out of free motor. Brain dynein was immobilised on the coverslip of a perfusion chamber by successively flowing reagents diluted in MRB80 buffer and incubating each for 10 minutes and washing with 30 μl MRB80 as follows: The coverslip was first activated by flowing in 0.1 mg ml^-1^ PLL(20)-g[3.5]-PEG(2)/PEG(3.4)-Biotin (50%) (#PLL(20)-g[3.5]-PEG(2)/PEGbi, Susos AG), followed by 0.625 mg ml^-1^ Streptavidin (#S4762 Sigma), a solution of 20 µg ml^-1^ biotinylated anti Mouse IgG antibody (#115-065-003-JIR-2ml STRATECH SCIENTIFIC LIMITED) and 20 µg ml^-1^ anti-Dynein Intermediate chain antibody (#MAB16818 Sigma-Aldrich). Unreacted streptavidin binding sites were then blocked using biotin-BSA (#PN29130, Fisher Scientific) and any unspecific sites blocked using 1 mg ml^-1^ κ-casein (#C0406 Sigma) solution. 30 nM dynein was then flown into the chamber and was incubated for 30 minutes. The chamber was washed with 30 μl image buffer (MRB80 supplemented with 0.6 mg ml ^-1^ κ-casein, 2 mM Mg-ATP, 4 mM DTT, 50 mM glucose (#G8270, Sigma), 0.2 mg ml ^-1^ catalase (#C9322, Sigma), and 0.4 mg ml ^-1^ glucose oxidase (#G7141, Sigma)). GMPCPP-stabilised HiLyte-647 labelled microtubules diluted 1:100 in image buffer were flown into the chamber. The chamber was sealed with wax to prevent evaporation and microtubule movement was recorded on an Olympus TIRF microscope using an 100X, NA 1.49 objective, 1.6X additional magnification, 60 mW 488 nm, 50 mW 561 nm, 100mW 640 nm laser lines, and a Hamamatsu ImageEM-1k back-illuminated EM-CCD camera (Hamamatsu Photonics) under control of xCellence software (Olympus). All experiments were carried out at 30 °C inside an incubator (Okolab, Ottaviano, Italy). Microscopy images were processed using ImageJ, and the length of all microtubules on the 1^st^ (0 minutes) and 600^th^ (40 minutes) frames of each stacks were measured using the line tool in ImageJ.

### Microtubule bundling assays

#### Dynein-mediated bundling in solution

GMPCPP-stabilised HiLyte-647 and biotin-labelled microtubules were incubated with 30nM recombinant Alexa 488 labelled human dynein in image buffer without κ-casein for 10-60 minutes at RT. Alongside this incubation, a perfusion chamber was prepared by flowing in PLL(20)-g[3.5]-PEG(2)/PEG(3.4)-Biotin (50%) and Streptavidin as described above. The chamber was then washed with 30 μl MRB80 followed by 10 μl image buffer without κ-casein. Then the reaction mix containing microtubules and dynein was introduced into the perfusion chamber. The chamber was sealed with wax and images recorded using the TIRF setup described above using the following settings: 100 ms exposure time, at a rate of 2 frames per second for 5 minutes or 4 s timelapse for 40 minutes, and 18% laser power for both 488 and 640 nm lasers. Microtubule bundles were distinguished from single microtubules due to their increased brightness. Single and bundled microtubules were marked with the line tool in ImageJ and kymographs for recording dynein motion were created in the 488 nm channel. To determine microtubule polarity within bundles, the number of dynein molecules moving from left to right and right to left was counted manually and the ratio of the larger of the two counts to total dynein molecules calculated as the microtubule polarity index. To determine the fraction of sliding bundles, kymographs were generated from the 640 nm channel and events counted where diagonal streaks were observed indicating movement of one of the microtubules in the bundle. The motility of dynein molecules on single microtubules and in bundles was analysed using a custom macro in ImageJ. Each molecule was manually traced in kymographs using the segmented line tool and phase length, phase speed, total dwell time, run length and average speed calculated for each molecule.

#### Splitting of microtubule bundles by kinesin

A perfusion chamber was activated by flowing in Drosophila kinesin-1 heavy chain in MRB80 and incubation for 30 minutes at RT. The chamber was then washed with 30 μl MRB80 and 10 μl Image buffer. Preformed GMPCPP stabilized HiLyte 647 microtubule bundles with Dynein (60 minutes incubation) was flown into the chamber. The chamber was sealed with wax and images were recorded in TIRF mode with the following imaging settings: Exposure time, 300 ms, 4 s timelapse for 5 minutes, and laser power 18 mW 640 nm laser as excitation. Number of microtubules per bundle was determined by counting the number of components microtubules, and polarity of the bundle was determined form the direction of the movement of component microtubules as they split from a bundle by Kinesin-1.

#### Microtubule sliding assay

A perfusion chamber was prepared with PLL(20)-g[3.5]-PEG(2)/PEG(3.4)-Biotin and Streptavidin as described above. Later, biotin-HiLyte-647 labelled track microtubules in MRB80 were flown into the chamber. This was done to immobilise microtubule tracks on the surface. Unbound microtubules were washed out with 30 μl MRB80 after 5 minutes incubation. The surface was blocked using 1 mg ml^-1^ κ-casein solution in MRB80 for 5 minutes. The chamber was then washed with 10 μl image buffer. microtubule overlaps were formed by flowing in X-rhodamine-labelled transport microtubules together with 30 nM Alexa-488-labelled recombinant Dynein solution in Image buffer. The chamber was then sealed with wax and images were recorded in TIRF mode using the following settings. Exposure time, 300 ms, 4 s timelapse for 5 minutes, and laser power 18 mW for 488, 561, and 640 nm lasers.

### Force measurements

A perfusion chamber was prepared by sandwiching a cover glass (BDH, 22 × 32 mm #1) on top of acid and plasma cleaned cover glass (VWR 24 × 50mm #1.5) containing two lines of Corning High Vacuum Grease. The flow cell had a resulting volume of ∼10 µl. It was then activated by flowing in PLL(20)-g[3.5]-PEG(2)/PEG(3.4)-biotin followed by streptavidin as described above. Microtubules were bundled by incubating biotin-HiLyte-647 labelled microtubules together with 0.1 µM dynein solution in image buffer for 60 minutes at RT. These pre-bundled microtubules were then flown into the chamber and incubated for 10 minutes to allow binding of microtubules (both single and bundles) to the surface. A bead motor mix was prepared by mixing 560 nm plain polystyrene beads with dynein in buffer (80 mM PIPES, pH 7, 2 mM MgSO_4_, 1 mM EGTA, 1 mM DTT, 0.1 mg ml^-1^ β-casein, and 1 µM Mg-ATP) and incubated on ice for 10 minutes. The bead motor mix was further diluted 50 times in the assay buffer (MRB80, 1 mM Mg-ATP, 0.2 mg ml^-1^ κ-casein, 10 µM Taxol, 0.2 mg ml^-1^ catalase, 0.4 mg ml^-1^ glucose oxidase, 50 mM glucose, 1% glycerol) and was then flown into the chamber. The chamber was then placed onto the custom-built optical trapping microscope described previously [41]. Microtubules were visualised using in-built differential interference contrast (DIC). Individual beads were trapped using a 3 W Nd:YAG12 1064 nm laser (I E Optomech Ltd, Newnham, England) and positioned in the proximity of microtubules by steering the trap. Force recording was started by switching the imaging channel to 16 amplitude contrast and projecting the shadow of the bead onto the quadrant photodiode detector. Displacements data were acquired at RT, 20 kHz frequency, trap stiffness 0.013-0.055 pN nm^-1^. Typically, a recording lasted 180 s. A calibration for trap stiffness was done each day before commencement of measurements. Measurements were done under conditions when 25 % or less beads were running on microtubule bundles. Individual force traces were analysed by a custom R code (R core team, 2017). Stall force and the duration of runs were obtained from 100 ms moving average smoothened raw data file.. Runs were defined as events that reached a constant stall force (± 10 % fluctuation) for at least 100 ms before a sudden drop to baseline of at least 24 nm and manually marked. Force-velocity relationship was determined from manually identified runs using a custom R code determining the time taken to increase force in 0.5 pN increments until stall force is reached.

### Figure preparation and statistical analysis

Origin, R and MATLAB were used to create plots and perform statistical analysis. Statistical tests are indicated in corresponding figure legends, p values are either stated in the figure panel or indicated with asterisks and explained in the figure caption. Images and kymographs were prepared using ImageJ, manipulations were limited to adjusting minimum and maximum grey values, application of false-coloured and inverted look up tables and cropping of images. All schematics were prepared, and figures assembled using Adobe Illustrator. Data in the text are given as mean ± standard error of the mean (SEM).

**Supplementary Table 1:**
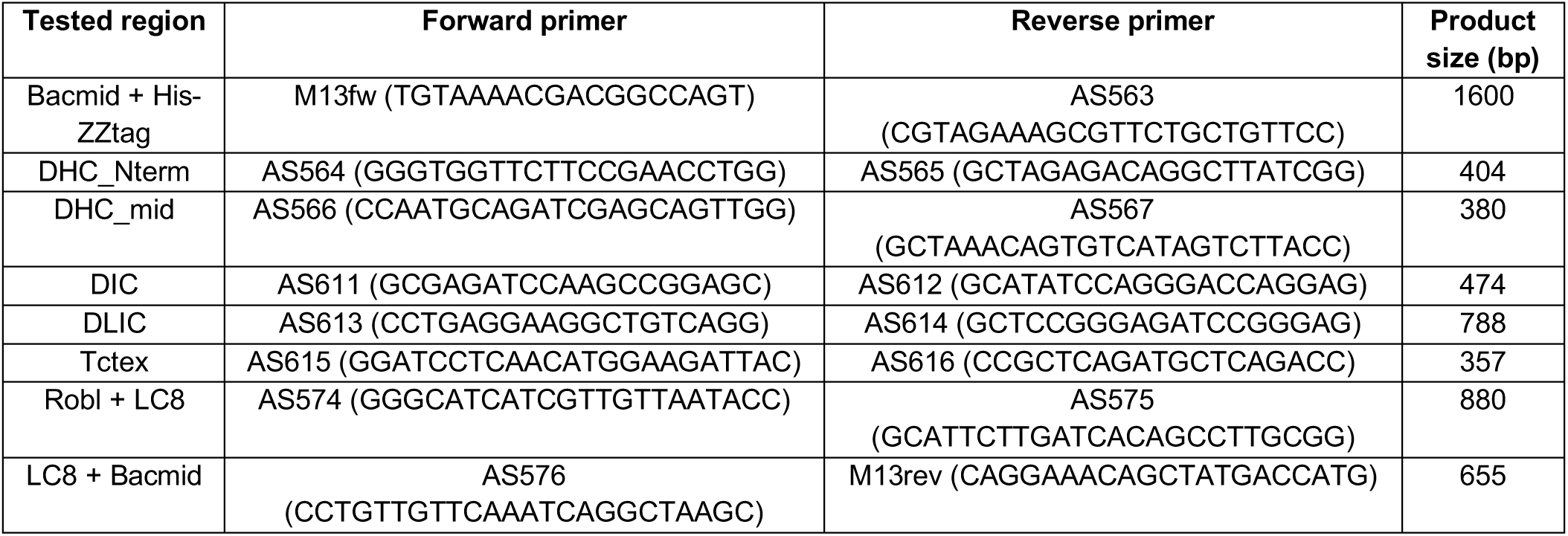
List of primers for PCR-based verification of complete pDyn3 integration.

